# Comparison of genomic assembly and annotation based on two clones of avian pathogenic *Escherichia coli*

**DOI:** 10.1101/2024.11.22.624809

**Authors:** Yufei Zhao, John Elmerdahl Olsen, Louise Poulsen, Henrik Christensen

## Abstract

Methods for assembly and annotation of whole genomic sequences were compared for six strains of avian pathogenic *Escherichia coli* (APEC). Two vertically transferred *E. coli* clones, represented by three isolates all belonging to pulse field genome electrophoresis (PFGE) type 65- sequence type (ST)95 and three isolates belonging to PFGE type 47- ST131, were selected for Illumina short read sequencing. There was no significant difference between SPAdes and CLC Genomic Workbench for benchmark parameters to assemble the short reads. The six strains were also sequenced by long read sequencing (Nanopore) and these reads were hybrid assembled with the short reads. Unicycler provided a lower number of contigs and higher N50 compared to Flye. No significant differences between total length of genomes were obtained from the four assemblers. At least 2.1 and 0.9% of coding gene sequences (CDSs) annotated with RAST and PROKKA, respectively were wrongly annotated. The errors were most often associated to CDS of shorter length (< 150 nt) with functions such as transposases, mobile genetic elements or being hypothetical. The investigation points out the importance of controlling automatic annotations and suggest further work to improve annotations in strains not belonging to the K12 or B lineages.

## 1. Introduction

High quality whole genomic sequences are needed for accurate annotation, identification of plasmids, to trace bacterial clones and more general to provide the basic information for modern investigation of bacteriology. The Illumina sequencing platforms can generate high accuracy (>99.99%) raw reads with a relatively low cost [1] and enable extraction of information of multi locus sequence type (MLST), prediction of virulence associated genes (VAGs), antibiotic resistance genes (ARGs) and provide clonal comparison by single nucleotide polymorphism (SNP) analysis [2]. However, the pair-end raw reads generated are often not longer than 250 to 300 base pairs, which makes it difficult to fully close a bacterial genome leaving several incomplete contiguous sequences (contigs). The fragmented contigs may hinder the determination of whether VAGs and ARGs are located on chromosomes or mobile genetic elements. Another problem is the lack of knowledge of contig orientation and orientation of annotation of structural repetitive regions and larger rearrangement such as transposons, insertion sequences and genetic cassette. The repeated sequences are often larger than the sequencing length of the Illumina reads making assembly impossible [1, 3]. The long read sequencing platforms, such as Pacific Biosciences (PacBio) and Oxford Nanopore Technologies (ONT) can generate single DNA reads with a median length of 10k bp and up to 100k bp [1, 4], and thus can be a great improvement to correct sequences generated with Illumina technology with respect to the repetitive and other complex regions on genomes [5]. Although the raw read accuracy can go up to 99.5% with the new R10.4.1 flow cell and V14 kit according to ONT (https://nanoporetech.com/), hybrid assemblies with a combination of the accuracy of short read sequence and the completeness of long read sequencing still provide the most reliable and cost-effective approach to reconstruct bacterial genomes [6, 7, 8).

Two of the most common pipelines for annotation are RAST [9, 10] and Prokka [11]. RAST predicts open reading frames (ORFs) based on GLIMMER 3.0 [12], combined with protein families “subsystems” as well as other tools and iterations. GLIMMER 3.0 is based on interpolated Markov model (IMM) search where ORFs are identified in reverse from the stop codon and back towards the start codon. Further optimization is by optimization for ribosomal binding sites using a position weight matrix improving the prediction of translation initiation sites. Prokka [11] uses Prodigal [13] to identify ORFs based on a dynamic programming approach optimizing start and start and stop codon identification using prediction of ribosome binding sites, GC contents, hexamer distributions as well as iterations of the process.

The aim of the study was to investigate the influence of assembly and annotation methods (RAST, Prokka) using *E. coli* strains of the avain pathogenic type (APEC) as model organisms. The comparison of closely related strains allowed the identification of annotation errors associated with both RAST and Prokka.

## 2. Material and methods

### 2.1 Sample collection and molecular typing

Six APEC strains were chosen for the current study. They represented two APEC clones of sequence type (ST) 95, pulse field gel electrophoresis type 65 and ST131, PFGE type (47), respectively [14] (Table 1).

**Table 1.** Strains investigated, genome length, genetic elements from six hybrid assembled genomes.

|  |  |  |  | Hybrid assembly |  | INDSC acc. no. |  |  |
| --- | --- | --- | --- | --- | --- | --- | --- | --- |
| Strain | Origin | Organ/lesion | PFGE-ST | Contig number | Contig length (bp) | Hybrid assembly | Short read assembly | Matching genomic elements |
| 729_021014_1_4 | Broiler parent | Salpingitis | 65-95 | 1 | 5,232,657 | CP160125 | JBELNO01 | Chromosome |
|  |  |  |  | 2 | 193,604 | CP160126 |  | Plasmid <sup>a</sup> |
|  |  |  |  | 3 | 6655 | CP160127 |  | Plasmid |
|  |  |  |  | 4 | 2676 | CP160128 |  | Plasmid |
|  |  |  |  | 5 | 2312 | CP160129 |  | Plasmid |
| 729_270514_x_12 | Broiler parent | Salpingitis | 65-95 | 1 | 5,232,714 | CP160121 | JBELNS01 | Chromosome |
|  |  |  |  | 2 | 193,604 | CP160122 |  | Plasmid <sup>a</sup> |
|  |  |  |  | 3 | 6655 | CP160123 |  | Plasmid |
|  |  |  |  | 4 | 2677 | CP160124 |  | Plasmid |
| 729_xx_1_3_yolk | Broiler | Yolk sac infection | 65-95 | 1 | 5,232,647 | CP160116 | JBELNM01 | Chromosome |
|  |  |  |  | 2 | 193,604 | CP160117 |  | Plasmid <sup>a</sup> |
|  |  |  |  | 3 | 6655 | CP160118 |  | Plasmid |
|  |  |  |  | 4 | 2676 | CP160119 |  | Plasmid |
|  |  |  |  | 5 | 2312 | CP160120 |  | Plasmid |
| 723_010814_2_1 | Broiler parent | Salpingitis | 47-131 | 1 | 4,866,047 | CP160112 | JBELNC01 (SSQU01) | Chromosome |
|  |  |  |  | 2 | 201,035 | CP160113 |  | Plasmid <sup>a</sup> |
|  |  |  |  | 3 | 6668 | CP160114 |  | Plasmid |
|  |  |  |  | 4 | 1552 | CP160115 |  | Plasmid |
| 729_141114_1_15 | Broiler parent | Salpingitis | 47-131 | 1 | 4,887,791 | CP160108 | JBELNB01 | Chromosome |
|  |  |  |  | 2 | 201,054 | CP160109 |  | Plasmid <sup>a</sup> |
|  |  |  |  | 3 | 100,533 | CP160110 |  | Plasmid |
|  |  |  |  | 4 | 1552 | CP160111 |  | Plasmid |
| 723_220514_1_6 | Broiler | Liver/septicaemia | 47-131 | 1 | 4,866,598 | CP160104 | JBELNH01 | Chromosome |
|  |  |  |  | 2 | 200,992 | CP160105 |  | Plasmid <sup>a</sup> |
|  |  |  |  | 3 | 6668 | CP160106 |  | Plasmid |
|  |  |  |  | 4 | 1552 | CP160107 |  | Plasmid |

<sup>a</sup> ColV-like

### 2.2 DNA extraction

*E. coli* isolates were cultured from stocks stored frozen in glycerol at - 80 °C. Cultivation was initiated on blood agar (Tryptose blood agar base Difco, Brøndby, Denmark) added 5% bovine blood with subsequent incubation at 37 °C for 24 h. A single colony from each plate was cultured in Luria-Bertani Broth (LB) at 37 °C overnight. The DNA was extracted with DNeasy Blood&Tissue Kit (QIAGEN) and Maxwell RSC Cultured Cells DNA kit (Promega, Madison, USA) according to manufacturer’s instructions. The purity and concentration of extracted DNA were evaluated by Nanodrop spectrophotometer (Thermo Fisher Scientific) and Qubit fluorometer (Thermo Fisher Scientific). The integrity of DNA was determined by gel-electrophoresis. DNA extracted from the isolates was used for both Illumina and ONT sequencing.

### 2.3 Library preparation and sequencing

DNA Libraries for Illumina sequencing were prepared using Illumina DNA Prep (Illumina, SanDiego, CA) and sequence reactions were prepared with Miseq Reagent kit v2 (Illumina, SanDiego, CA). Samples were sequenced by the Illumina MiSeq platform using 2 × 250 bp paired end sequencing strategy. In parallel, strains were sequenced by ONT on a MinION Mk1B sequencer (ONTechnology, Oxford). Libraries of selected DNA samples were prepared by using Rapid Barcoding Kit 96 V14 (SQK-RBK114, ONT), and were loaded on one MinION Flow Cell (R.10.4.1 FLO-MIN114, ONT) for sequencing for 48 h with an accurate model (260 bps). MinKNOW software v5.4.3 was used to perform the sequencing and data acquisition. Post-run base-calling and demultiplexing were performed with Guppy v6.4.8 (https://nanoporetech.com/document/Guppy-protocol, accessed 20 February 2025).

### 2.4 Raw reads quality control and trimming

Quality control, adapter removal and short reads filtering of Illumina reads were performed on fastp v0.12.4 for Linux with default parameter settings [15]. For ONT long reads, the short and low-quality reads were filtered by Filtlong v0.2.1 (https://github.com/rrwick/Filtlong) and quality control of long reads were assessed using SeqKit v2.5.1 [16].

### 2.5 Genome assembly and annotation

SPAdes v3.13.1 was used with a ‘--careful’ parameter for genome assemblies [17] for short reads generated by Illumina. Contigs smaller than 1000 bp were removed from the assembled genomes. CLC Genomic Workbench 22 (QIAGEN) was used for short reads assemblies to compare the assembly quality with SPAdes. Hybrid assemblies were performed by two workflows: Unicycler v0.5.0 was used directly with Illumina short reads and corresponding Nanopore long reads [4, 18]. The other approach used Flye v2.9.2 [19] to do a long-read assembly first, then the draft assemblies were polished with medaka v1.8.1 [20] and further improved with short reads by Pilon v1.2.2 [21]. Quast v5.0.2 [22] (Gurevich et al. 2013) and BUSCO v5.5.0 [23] were used to evaluate the quality of assemblies. Annotation was performed with Prokka v1.14.6 with ‘--force --centre X –compliant’ parameters [11] and the online toolbox Rapid Annotation using Subsystem Technology (RAST) [10].

### 2.6 Comparison of annotations

To control the differences among annotations, gene presence and absence matrices were established for RAST using Seed Viewer, and for Prokka using Roary. A Python script was written by chatGPT and used to parse the differential genes from matrices and extract corresponding sequences from original FASTA files. The sequences of differential genes were integrated into one Excel file. The sequences were aligned by BLASTn against the whole genome sequence of strains which did not annotate these genes within each clone. Different genes annotated with more than 95% identity were considered as annotation errors if they were annotated to the same DNA sequence. To illustrate selected regions, comparison of BLASTn alignments of differential region were performed using Clinker (https://cagecat.bioinformatics.nl/tools/clinker). The Venn diagrams showing relationships between Prokka and RAST annotations were generated on https://bioinformatics.psb.ugent.be/webtools/Venn/.

## 3. Results and discussion

### 3.1 Statistics of short read and hybrid assemblies

For the six strains, the average of read length of Illumina sequencing was 197.3 bp (range from 162.8 −262.4 bp) and 3909 bp (range from 2710 - 5129 bp) of ONT reads (Table 2). The average number of reads was 622,721 bp for Illumina and 109,933 bp for ONT. The coverage depth for all Illumina raw reads exceeded 30× with an average of 54×, while the ONT coverages ranged from 64× with an average coverage of 94× (Table 2). These parameters satisfy the common quality standards used for WGS assembly [24, 25, 26].

**Table 2.** Statistical analysis of reads.

| ST | Strain | Illumina |  |  |  | ONT |  |  |  |
| --- | --- | --- | --- | --- | --- | --- | --- | --- | --- |
|  |  | Mean read length (bp) | Number of reads | Total bases (Mbp) | Coverage (x)* | Mean read length (bp) | Number of reads | Total bases (Mbp) | Coverage (x) |
| 95 | 729_021014_1_4 | 184.9 | 515,309 | 154 | 41.1 | 5129 | 137,504 | 664 | 151.9 |
|  | 729_270514_x_12 | 190.5 | 613,533 | 186 | 50.3 | 4138 | 84,953 | 336 | 75.7 |
|  | 729_xx_1_3_yolk | 162.8 | 594,174 | 150 | 41.7 | 4260 | 95,064 | 387 | 87.2 |
| 131 | 723_010814_2_1 | 262.4 | 800,953 | 283 | 90.5 | 2710 | 109,803 | 283 | 64.1 |
|  | 729_141114_1_15 | 176.2 | 593,386 | 167 | 45.0 | 3832 | 138,333 | 508 | 114.2 |
|  | 723_220514_1_6 | 207.2 | 618,972 | 186 | 55.2 | 3386 | 93,939 | 301 | 68.5 |
|  | Average | 197.3 | 622,721 | 188 | 54.0 | 3909 | 109,933 | 413 | 93.6 |

\*coverage = (number of read × read length ) / total genome size. In which total genome size was estimated to 5.2 Mbase.

QUAST and BUSCO were used to test the assembly performance of the commonly used software CLC Genomic Workbench and SPAdes for short reads assembly and Flye, and Unicycler for hybrid assemblies (Table 3). The number of contigs did not differ significantly between CLC Genomic Workbench and SPAdes (Fisher’s exact test, P = 0.819). For hybrid assemblies, Unicycler provided lower number of contigs than Flye in both clones (*P* = 0.0003, Fisher’s exact test). NG50 is the shortest contig length of an assembly which is 50% of the reference genome or longer. N50 did not differ significantly between SPAdes and CLC (Fisher’s exact test, P = 0.768), however, longer N50 was obtained by Unicycler than from Flye (Fisher’s exact test, P = 0.0002). The total length of genomes did not differ significantly between CLC Genomics Workbench, SPAdes, Flye and Unicycler.

**Table 3.** The quality of assemblies generated by CLC Genomic Workbench and SPAdes for short-read only assemblies and Flye and Unicycler for hybrid assemblies.

| ST | Strain | Number of contigs |  |  |  | NG50 (bp) |  |  |  | Total length (bp) |  |  |  | Completeness (%) |  |  |  |
| --- | --- | --- | --- | --- | --- | --- | --- | --- | --- | --- | --- | --- | --- | --- | --- | --- | --- |
|  |  | CLC | SPAdes | Flye | Unicycler | CLC | SPAdes | Flye | Unicycler | CLC | SPAdes | Flye | Unicycler | CLC | SPAdes | Flye | Unicycler |
| 95 | 729_021014__1_4 | 110 | 99 | 5 | 5 | 169,077 | 196,246 | 5,232,681 | 5,232,657 | 5,310,368 | 5,312,171 | 5,444,670 | 5,437,904 | 100 | 100 | 100 | 99.8 |
|  | 729_270514_x_12 | 100 | 100 | 5 | 4 | 211,910 | 204,287 | 5,232,730 | 5,232,714 | 5,313,797 | 5,314,927 | 5,449,369 | 5,435,650 | 100 | 99.8 | 100 | 99.8 |
|  | 729_xx_1_3_yolk | 143 | 132 | 6 | 5 | 135,314 | 164,290 | 5,232,625 | 5,232,647 | 5,307,620 | 5,316,910 | 5,446,817 | 5,437,894 | 100 | 100 | 100 | 99.8 |
| Average |  | 117.7 | 110.3 | 5.3 | 4.7 | 172,100.3 | 188,274.3 | 5,232,678.7 | 5,232,672.7 | 5,310,595.0 | 5,314,669.3 | 5,446,952.0 | 5,437,149.3 | 100.0 | 99.9 | 100.0 | 99.8 |
| 131 | 723_010814_2_1 | 46 | 45 | 16 | 4 | 354,765 | 353,089 | 1,074,415 | 4,866,047 | 5,033,610 | 5,022,129 | 5,076,852 | 5,075,302 | 100 | 100 | 100 | 99.8 |
|  | 729_141114_1_15 | 88 | 88 | 5 | 4 | 192,789 | 224,060 | 4,949,514 | 4,887,791 | 5,110,669 | 5,115,410 | 5,164,053 | 5,190,930 | 100 | 99.8 | 100 | 99.8 |
|  | 723_220514_1_6 | 51 | 50 | 8 | 4 | 354,390 | 353,089 | 2,855,420 | 4,866,598 | 5,021,087 | 5,022,103 | 5,083,114 | 5,075,810 | 100 | 100 | 100 | 99.8 |
| Average |  | 61.7 | 61.0 | 9.7 | 4.0 | 300,648.0 | 310,079.3 | 2,959,783.0 | 4,873,478.7 | 5,055,122.0 | 5,053,214.0 | 5,108,006.3 | 5,114,014.0 | 100.0 | 99.9 | 100.0 | 99.8 |

Considering the performance of Unicycler in giving lower number of contigs and higher NG50 values, it was decided to be the best assembler for downstream annotation.

### 3.2 Comparison of annotations for hybrid assemblies

Annotations were compared for RAST and Prokka (Table 3). Identical number of rRNA genes (n = 22) were predicted by both annotation methods in all strains. The number of tRNA genes differed slightly. One tmRNA was annotated in all strains by Prokka compared to none by RAST. Different numbers of coding DNA sequences (CDS) were found for the hybrid genomes when annotated by RAST and Prokka. Overall, Prokka annotated less (187-327) CDSs compared to RAST (Table 4). RAST predicted that strains within clone PFGE65-ST95 differed by 88 CDS and strains of PFGE47-ST131 by 235 CDS. Meanwhile, Prokka predicted 47 different CDS in PFGE65-ST95 and 200 in PFGE47-ST131 (Fig. 1 A C).

**Fig. 1.**
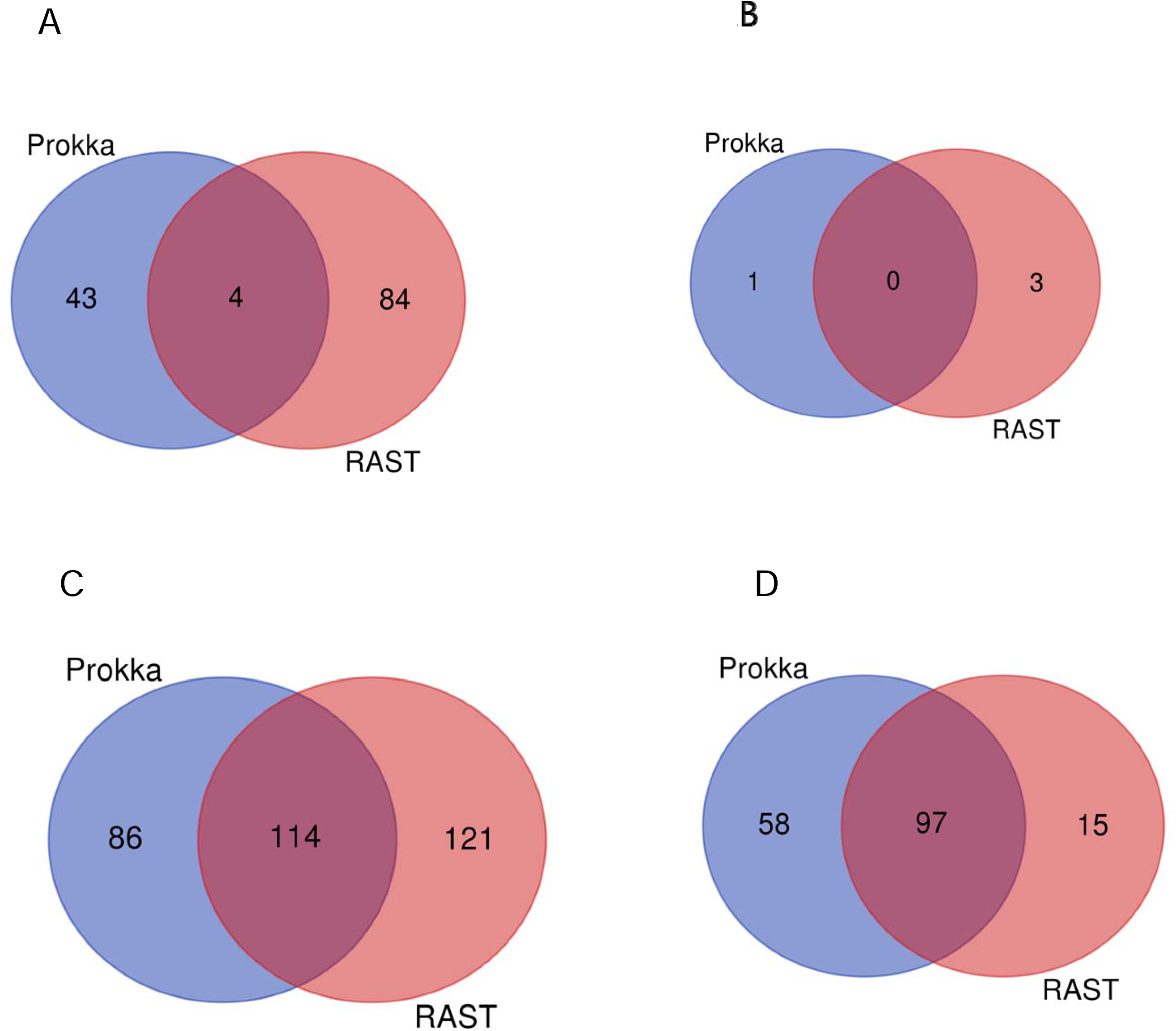
Venn diagrams illustrating the deviating annotations between Prokka and RAST. For PFGE65-ST95, (A) shows the number of different genes predicted before BLASTn control and (B) shows the numbers after control. For PFGE47-ST131, (C) shows the number of different genes predicted before BLASTn control and (D) shows the numbers after control.

**Table 4.** Statistical analysis of annotations.

| ST | Strain | CDS |  | tRNA |  | tmRNA |  | Total *RNA |  | Number of CDS differences |  |
| --- | --- | --- | --- | --- | --- | --- | --- | --- | --- | --- | --- |
|  |  | RAST | Prokka | RAST | Prokka | RAST | Prokka | RAST | Prokka | RAST | Prokka |
| 95 | 729_021014_1_4 | 5476 | 5149 | 90 | 92 | 0 | 1 | 112 | 115 | 88 | 47 |
|  | 729_270514_x_12 | 5450 | 5263 | 90 | 92 | 0 | 1 | 112 | 115 |  |  |
|  | 729_xx_1_3_yolk | 5442 | 5148 | 90 | 92 | 0 | 1 | 112 | 115 |  |  |
| 131 | 723_220514_1_6 | 4967 | 4699 | 84 | 86 | 0 | 1 | 106 | 109 | 235 | 200 |
|  | 723_010814_2_1 | 4958 | 4698 | 84 | 86 | 0 | 1 | 106 | 109 |  |  |
|  | 729_141114_1_15 | 5094 | 4830 | 85 | 86 | 0 | 1 | 107 | 109 |  |  |

\*22 rRNA genes in all strains for both annotation methods.

The reasons for the different numbers of CDS annotated by RAST and Prokka were analysed. Annotation comparison was performed based on presence and absence matrix generated from both RAST and Prokka. It was then surprising to find divergent annotations between strains for exactly the same DNA sequences. For RAST annotations of strains of the PFGE65-ST95 clone, 71 out of the 88 genes annotated differently between the three strains got 100% identity in complementary genomes, when controlled on sequence level showing that they were error prone annotations. Another 14 genes got more than 97% identity for at least one strain by BLASTn leaving three real gene differences in 729_270514_x_12 since the strain did not harbor the 2312 bp plasmid with three CDSs (Fig. 1 A B).

Similar problematic annotations were obtained among PFGE65-ST95 genomic sequences annotated by Prokka, in which 13 of 47 genes annotated differently within the three strains of PFGE65-ST95 got hits of 100% identity. Another 32 genes got more than 97% identity and one gene got 94.7% identity. These 46 genes were regarded as error prone annotations with Prokka (Fig. 1 A-B).

In PFGE47-ST131, with RAST annotation, 120 out of 235 genes annotated differently got 100% identity in whole genome sequences which were not annotated for these genes and three gene differences were identified the same way with more than 98% identity. All 123 annotations were considered error prone. Ninety-eigth real gene differences were related to contig 3 identified as a plasmid where 723_010814_2_1 and 723_220514_1_6 carried a 6668-bp *ColRNAI* related plasmid and 729_141114_1_15 carried a 100,533-bp Incl-1 plasmid. Another 14 real differences originated in phage related insertions into the chromosome of 729_141114_1_15 (Fig. 1 C – D).

For PFGE47-ST131 genomic sequences annotated with Prokka, 25 of 200 different genes got 100% identity in reciprocal sequences which were not annotated for these genes and 20 different genes got more than 97% identity. These 45 annotations were considered error prone. Similar to RAST annotation, 141 gene differences were caused by different genes on contig 3 (unique IncI-1 plasmid), and 14 annotation differences related to prophage regions in the chromosome of strain 729_141114_1_15 (Fig. 1 C – D).

Fig. 2 shows examples of inaccurately annotated genes and their flanking regions visualized by Clinker. The *neuO* gene (polysialic acid O-acetyltransferase) was only annotated in strain 729_270514_x_12 and not in the other two strains although they had identical DNA sequences in this region (A). In 729_141114_1_15, the duplicate annotation for the *pdeL* gene as *pdeL* and *pdeC* in strain 729_141114_1_15 compared to the other two strains was found to be an error since all three strains shared 100% sequence identity (B). The last example is with two copies of the transposase IS*Cep1* gene in two strains which were only annotated as one in 729_141114_1_15. Also here regions compared on the figure shared 100% sequence identity.

**Fig. 2.**
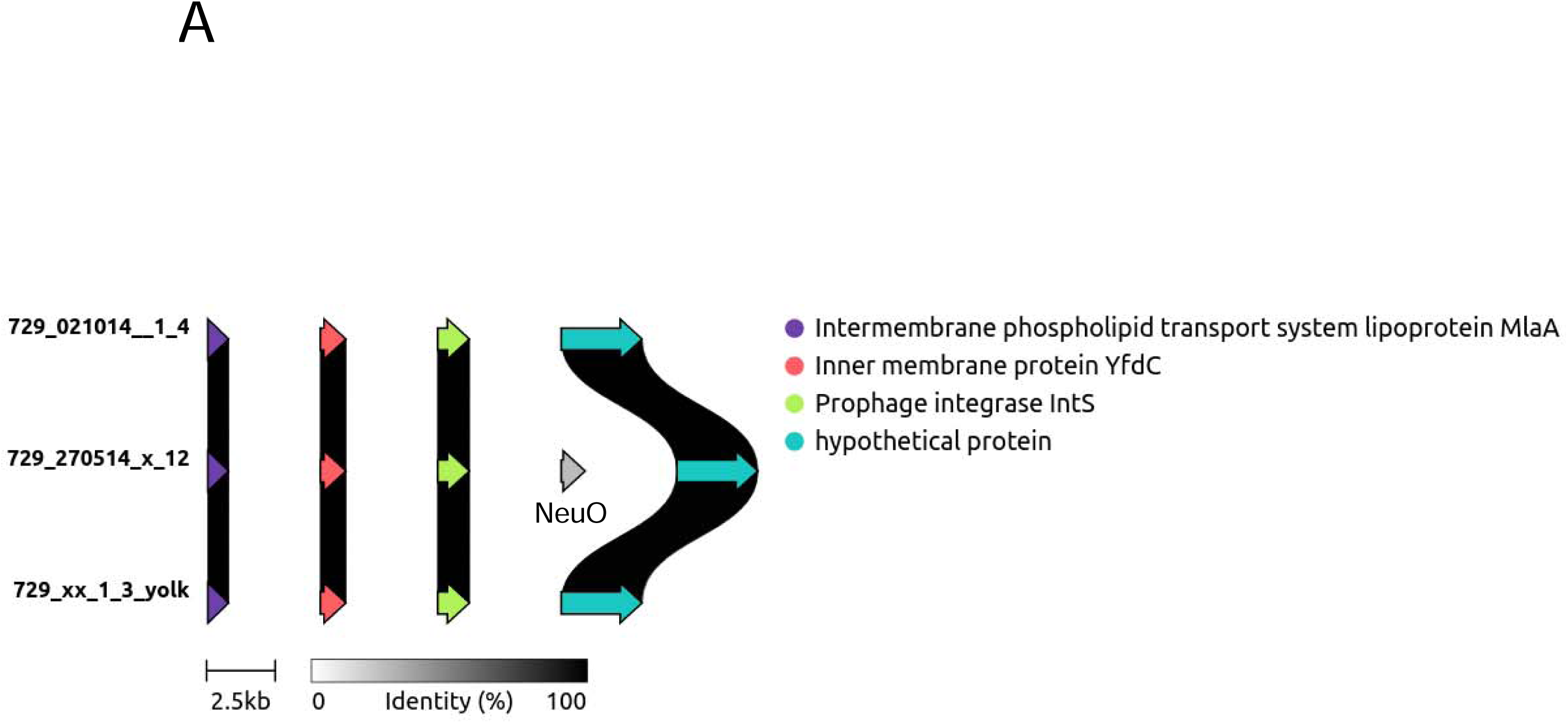
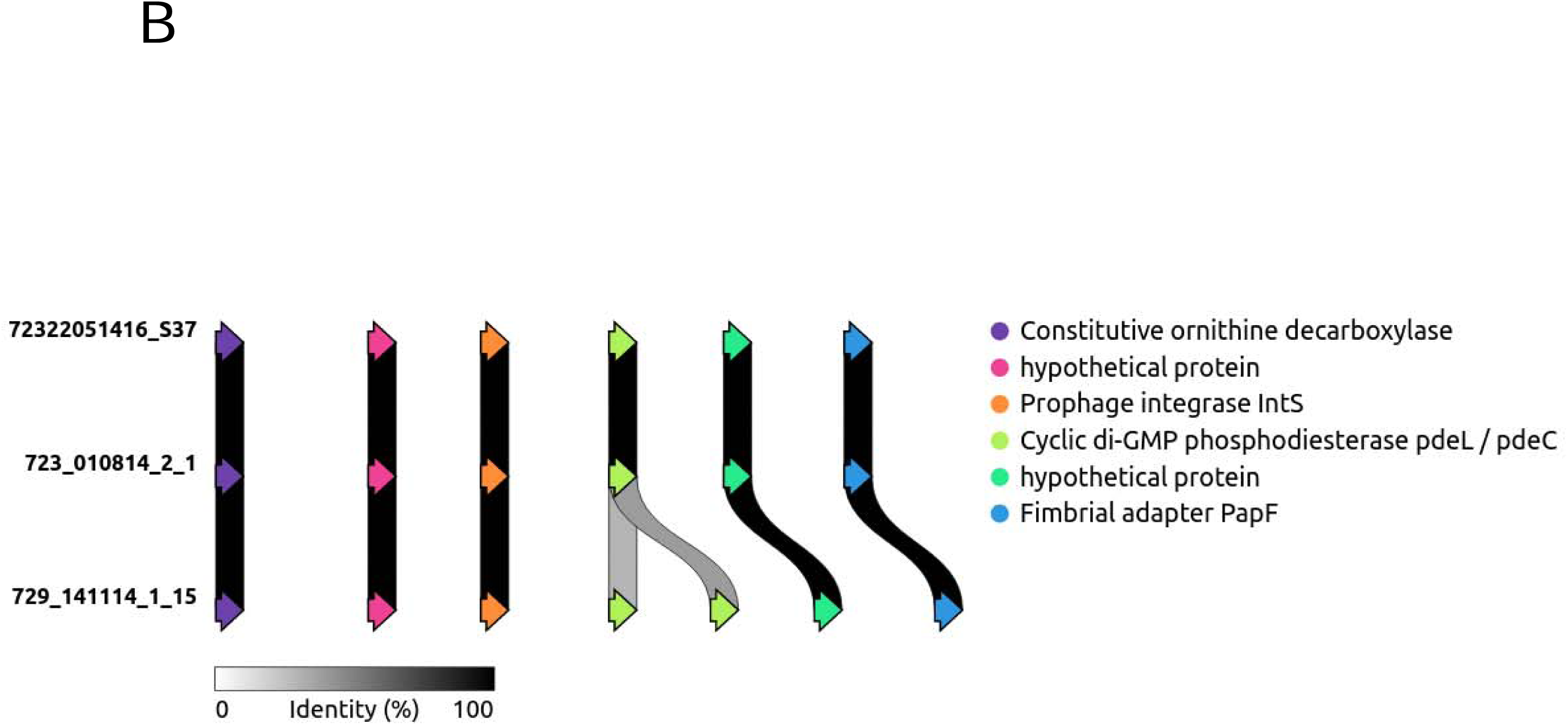
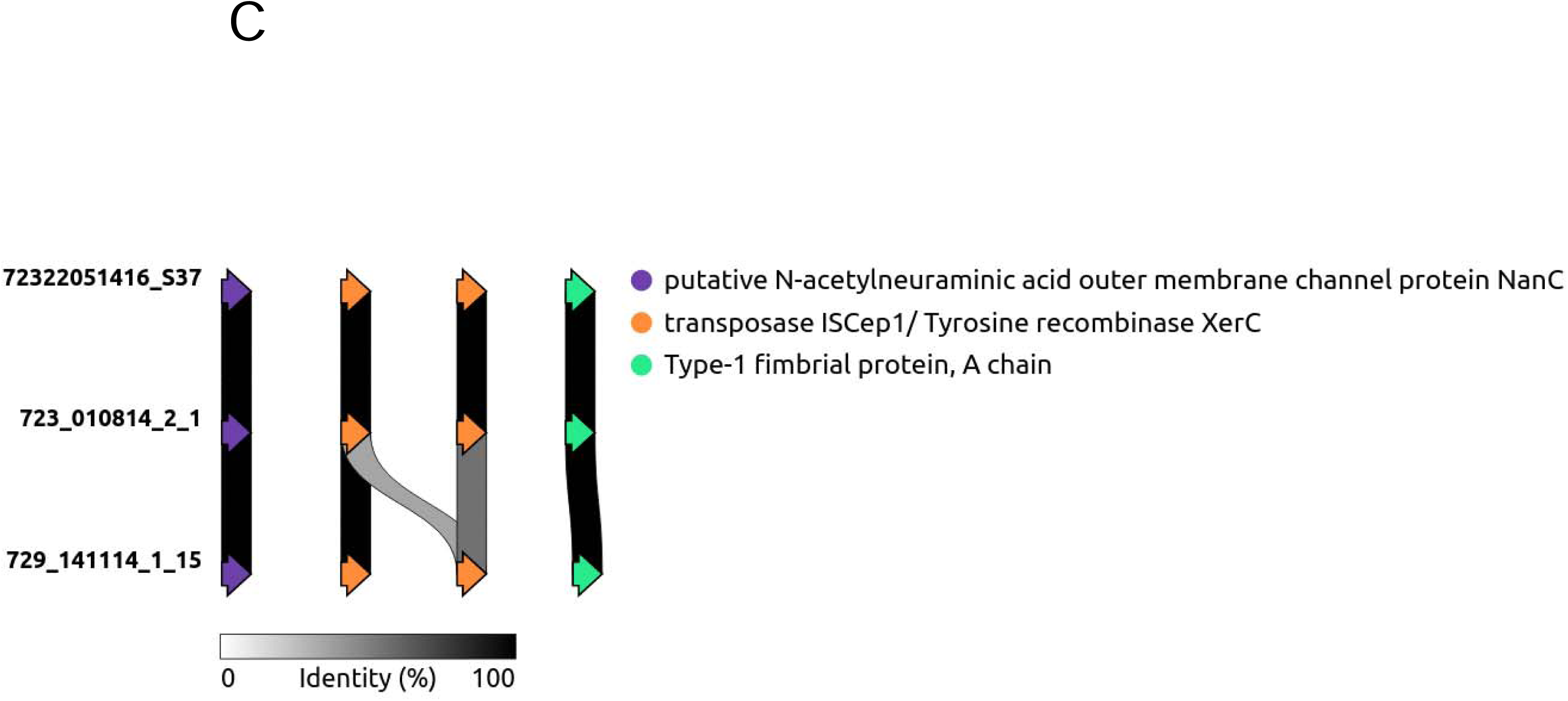
Examples of genetic comparisons of inaccurately annotated genes from Prokka and their flanking regions. An additional annotation is given for the *neuO* gene in 729_270514_x_12 (A). In 729_141114_1_15, duplicate annotation for the *pdeL* gene as *pdeL* and *pdeC* was found (B). Two copies of the transposase IS*Cep1* gene in two strains were only annotated as one in strain 729_141114_1_15 (C). The regions compared all shared 100% DNA sequence identity without gaps.

In all, 2.1 and 0.9% of CDSs annotated with RAST and PROKKA, respectively were error prone. The errors were most often associated to CDSs of shorter length (< 150 nt) with functions annotated as transposases, mobile genetic elements or hypothetical. These estimates are only based on reciprocal differences between the annotations and may be underestimated and not included CDSs wrongly annotated in all strains compared.

The divergence associated with annotations of identical DNA sequence regions falls back on the annotating methods. The methods and procedures in RAST and Prokka are quite different. RAST predicts ORFs based on GLIMMER 3.0 where ORFs are predicted by IMM search in reverse from the stop codon and back towards the start codon followed by optimization for ribosomal binding sites improving the prediction of translation initiation sites. Optimizations from previous versions of the program included in particular the prediction of translation initiation sites and one reason for inaccurate ORF prediction can be that further optimization of this step is needed. Prokka uses Prodigal [13] to identify ORFs based on a dynamic programming approach optimizing start and start and stop codon identification by prediction of ribosome binding sites, GC contents, hexamer distributions as well as iterations of the process. A lower limit of 90 bp is used in the program to score start-stop codon pairs [13] and this may be some of the reason for the errors observed for shorter genes in the current study. Another reason for errors with Prodigal may be the use of a window of 5 kb to identify start and stop codons. In the current comparison it can be anticipated that sequence divergence between two strains in such a window may lead to different annotations at local level even though sequenced are identical for this region. Hyatt et al. [13] found that Glimmer 3 predicted around 4% more ORFs in E. coli K12 compared to Prodigal which is in line with the current study. A problem when comparing annotations is the lack of a golden standard. Such a standard could have been obtained from *E. coli* strain K12 variant MG1655 where annotation has been refined for decades for instance by EcoGene [27, 28]. Unfortunately this resource seems no longer maintained and accessible. Considering the high genomic variation between reference and field strain of *E. coli* a full comparison would not have been possible. Further optimization of the search parameters in the annotation programs may be needed and targeted better towards *E. coli* and further curation of annotations needed.

## 4. Conclusion

There was no significant difference between SPAdes and CLC Genomic Workbench for benchmark parameters to assemble short reads. However, for hybrid assemblies, Unicycler provided lower number of contigs and higher N50 than Flye. In the comparisons, at least 2.1 and 0.9% of genes annotated with RAST and PROKKA, respectively were false annotations stressing the importance of controlling the annotations. This investigation showed that strains with closely related genomes at DNA level obtained differences in annotation for local regions with identical DNA sequences. The problem was identified with the most commonly used annotation programs RAST and Prokka. The errors are related to the statistical methods used for prediction of ORFs where differences in up or downstream regions may influence the prediction.

## CRediT authorship contribution statement

Yufei Zhao: Formal analysis, Investigation, Writing – original draft. Henrik Christensen: Strains, Formal analysis, Investigation, Writing – original draft. Louise Ladefoged Poulsen: Strains, Writing – review & editing. John Elmerdahl Olsen: Investigation, Resources, Conceptualization, Writing – review & editing, Supervision.

## Conflicts of interest

The authors declare that there are no conflicts of interest.

## Funding information

This study was partially funded by a State Scholarship Fund to Yufei Zhao from China Scholarship Council (CSC No. 202206350041).

## Repositories

GenBank under accession numbers in Table 1.

## Acknowledgements

Xiao Fei and Yaovi Mahuton Gildas Hounmanou are thanked for help with the analysis of Nanopore sequencing data.

## Notes

### Competing Interest Statement

The authors have declared no competing interest.

### Summary of Updates

Formatted with numbered sections, references in numbered format, extension of conclusion, CRediT statement

